# Systemic translocation of *Staphylococcus aureus* promotes autoimmunity: implications in autoantibody-mediated poor immune reconstitution from antiretroviral therapy in HIV

**DOI:** 10.1101/2025.08.04.668434

**Authors:** Da Cheng, Zhenwu Luo, Wangbin Ning, Sonya L. Heath, Magnus Gisslen, Richard W Price, Ruth Adekunle, Tabinda Salman, Douglas Johnson, John E McKinnon, Lishomwa C. Ndhlovu, Reafa Hossain, Wenhui Hu, Wei Jiang

## Abstract

**Background:** In 2017, our group first demonstrated that autoimmunity contributes to HIV pathogenesis, even without autoimmune disease. This concept is now broadly recognized, exemplified by the role of autoimmunity in severe COVID-19. In people with HIV (PWH) on suppressive ART, anti-CD4 autoantibodies may impair CD4+ T cell recovery, though the mechanisms driving their production remain unclear. Building on evidence from our group and others that *Staphylococcus aureus* and its peptidoglycan (PGN) promote autoimmunity, we investigated their contribution to anti-CD4 IgG in HIV.

**Methods:** Plasma from 32 ART-naive PWH, 53 ART-treated PWH, and 32 HIV-negative controls was analyzed for IgG autoantibodies and markers of *S. aureus* translocation using protein array, ELISA, and microarray. EcoHIV mice were injected intraperitoneally with saline, *S. aureus* PGN, or *Bacillus subtilis* PGN. PGN structures were compared by mass spectrometry.

**Results:** Among 87 autoantibodies, 40% were elevated in ART-naive PWH and largely normalized by ART; however, anti-CD4 IgGs remained elevated in PWH on ART. Anti-CD4 IgG levels inversely correlated with CD4+ T cell counts in ART-treated PWH and positively with *S. aureus* translocation. In mice, *S. aureus* PGN induced anti-CD4 IgGs, reduced gut CD4+ T cells, and promoted surface IgG binding and apoptosis in CD4+ T cells.

**Conclusion:** *S. aureus* and its PGN translocation may drive anti-CD4 autoimmunity and hinder immune recovery in PWH on suppressive ART, highlighting *S. aureus* colonization as a therapeutic target and supporting the development of competitive probiotic interventions such as *Bacillus subtilis*.

## Introduction

Recent studies, including ours, show that autoimmunity contributes to pathogenesis in infectious diseases without clinical autoimmune conditions (1). About 20% of PWH on ART fail to restore CD4+ T cell counts despite viral suppression, increasing comorbidity and mortality risks (2). We demonstrated that anti-CD4 autoantibodies drive CD4+ T cell death via ADCC, contributing to poor immune reconstitution post-ART (3), a finding supported by other researchers (4), though outcomes vary (5). The mechanisms driving anti-CD4 IgG production in HIV remain unclear.

Our prior work (6) demonstrated that S. aureus PGN induces pathogenic autoantibodies and kidney injury in SLE. Intraperitoneal injection of heat-killed S. aureus or its PGN elicited autoantibody responses in both C57BL/6J and autoimmune-prone mice (6, 7). As a major antigenic cell wall component, PGN exerts distinct immune effects depending on its bacterial source (8). Because S. aureus colonization is more frequent in PWH (9), we examined links between autoantibodies and CD4+ T cell counts and tested the role of S. aureus PGN in anti-CD4 IgG production. Our findings show that S. aureus PGN promotes anti-CD4 IgG generation, leading to CD4+ T cell loss and impaired immune recovery in PWH on suppressive ART.

## Results

### Autoantibody landscape and the unique characteristics of anti-CD4 IgGs

Among the 87 autoantibody IgGs (autoIgGs) in HIV+/ART+, 35 autoIgGs (40%) were elevated, 3 were reduced, and the rest were similar to controls (Figure 1A). ART normalized most autoIgGs, but anti-CD4 and anti-prothrombin IgGs remained elevated in HIV+/ART+ (Figure 1A-1B). Anti-CD8 IgG was lower in HIV+/ART-naive versus controls (Figure 1C). Plasma anti-CD4 IgG levels inversely correlated with CD4+ T cell counts in HIV+/ART+ only (Figure 1D). Anti-CD4 IgM and IgA levels were comparable across groups. In HIV+/ART+, anti-CD4 IgG1 and IgG3, but not IgG2 or IgG4, were elevated (Figure S1A-S1B) (3, 10). IgG1 and IgG3 in humans support ADCC (11, 12). In addition, plasma levels of anti-prothrombin IgG were directly correlated with anti-CD4 IgGs (r = 0.22, P = 0.01), but not correlated with CD4+ T cell counts (r = -0.16, P = 0.22).

**Figure 1.**
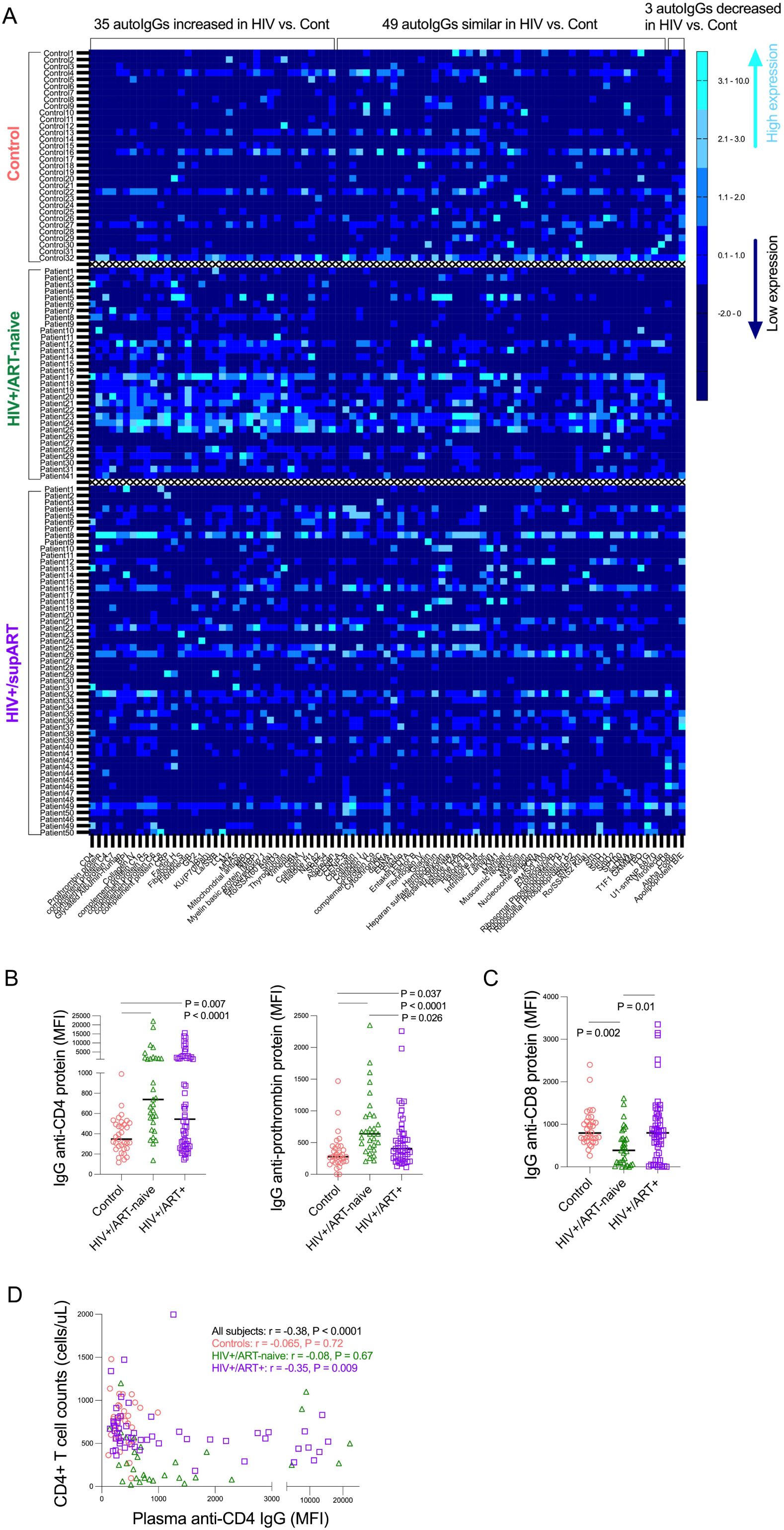
HIV-associated IgG autoantibodies. (A) Z-scored plasma levels of 87 autoantibodies in controls (n = 32), untreated PWH (n = 32), and ART-suppressed PWH (n = 53). (B) Anti-CD4 and anti-prothrombin IgG. (C) Anti-CD8 IgG. (D) Correlation between anti-CD4 IgG and CD4+ T cell counts. ANOVA and Spearman correlation. Samples were analyzed for autoantibodies using a protein array and collected from four institutions: Sahlgrenska University Hospital, San Francisco General Hospital, RHJ VAHCS, and MUSC.

### Elevated anti-CD4 IgG levels are associated with plasma levels of *S. aureus* translocation in PWH on suppressive ART

Our previous study in systemic lupus erythematosus (SLE) demonstrated that *Staphylococcus aureus* contributes to the production of pathogenic lupus-associated anti–double-stranded DNA IgG antibodies and to kidney damage (6). Based on these findings, we evaluated the potential role of *S. aureus* in the generation of anti-CD4 autoantibodies in HIV infection. To assess systemic exposure to *S. aureus*, we measured plasma IgG levels against *S. aureus* antigens as a surrogate of bacterial translocation. We found that plasma IgG levels against *S. aureus* antigens were positively correlated with anti-CD4 IgG levels in HIV+/ART+ individuals. (Figure 2A-2B).

**Figure 2.**
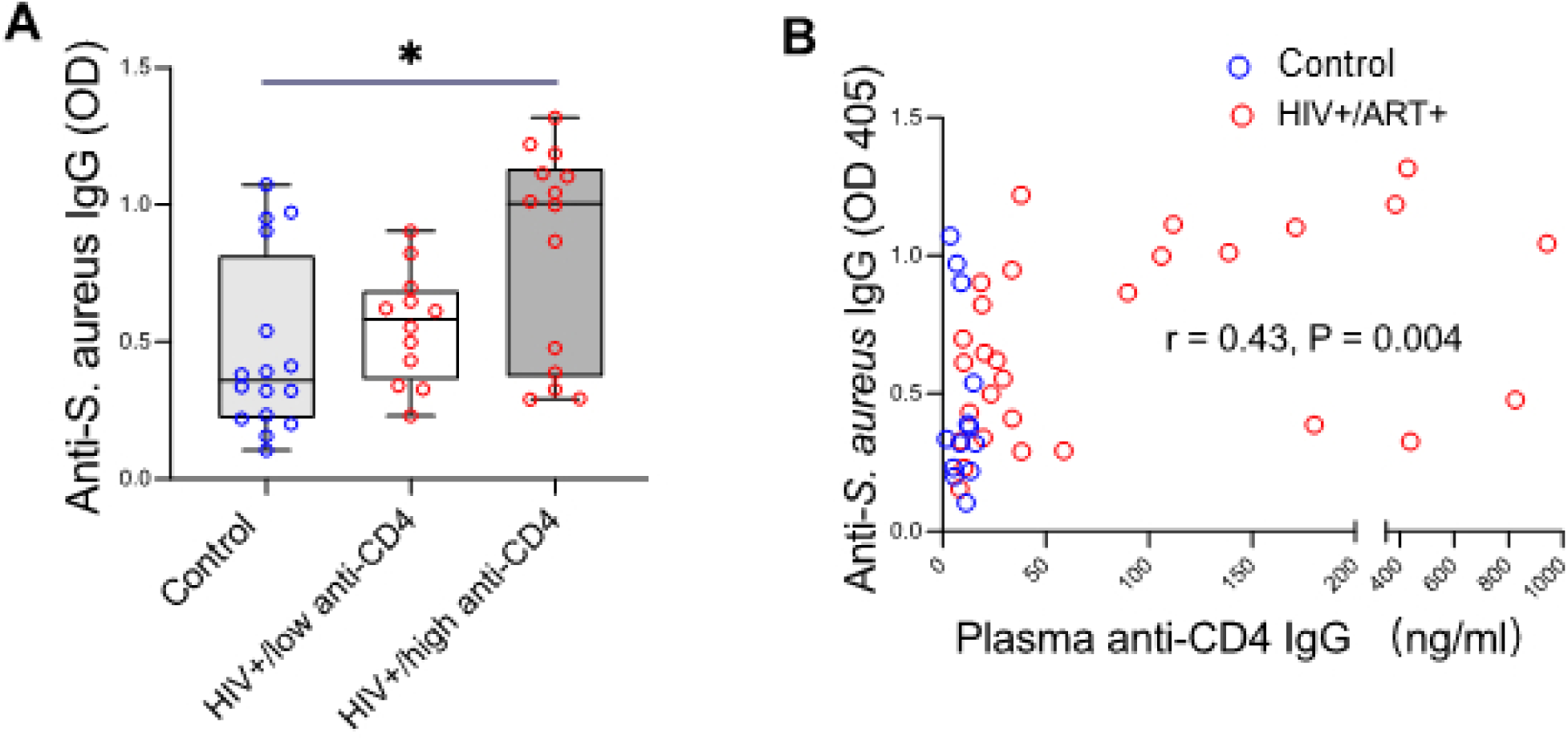
Association of S. aureus markers with anti-CD4 IgG in PWH on ART. (A) Correlation of plasma anti-CD4 IgG with anti-S. aureus IgG. (B) Elevated plasma anti-S. aureus IgG in HIV+/ART+. Spearman correlation and ANOVA. PWH were divided into low and high anti-CD4 IgG subgroups using a 35 ng/ml cutoff, based on the 95^th^ percentile in controls. Samples from MUSC were analyzed for anti-CD4 IgG and anti-S. aureus IgG by ELISA.

### *S. aureus* PGN induced anti-CD4 IgG production in EcoHIV mice

EcoHIV mouse study design is shown in Figure 3A and infection was verified by qPCR (Figure S2). S. aureus PGN increased serum anti-CD4 IgG without affecting total IgG (Figure 3B), indicating limited polyclonal B cell activation. In mice, IgG2a and IgG2b mediate ADCC (13), and S. aureus PGN selectively increased anti-CD4 IgG2b and IgG3 (Figure 3C). By contrast, B. subtilis PGN, a probiotic (14), did not induce anti-CD4 IgG (Figure 3B–3C).

**Figure 3.**
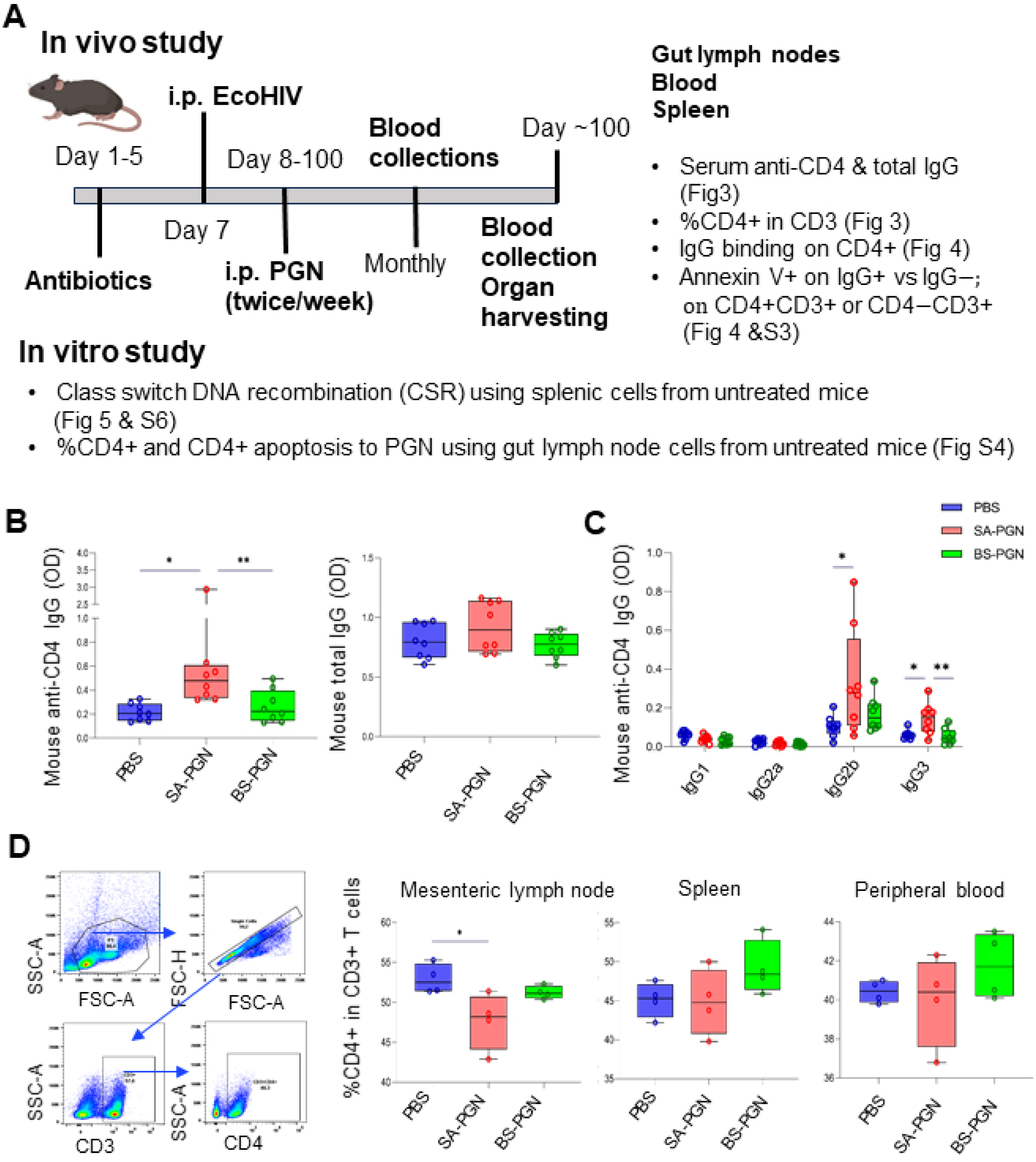
S. aureus PGN induces anti-CD4 IgG and reduces gut CD4+ T cells in EcoHIV mice. C57BL/6J mice received i.p. PBS, S. aureus PGN, or B. subtilis PGN. (A) Mouse study design. (B–C) Serum anti-CD4 IgG, total IgG, and IgG subclasses. (D) Gating strategy and CD4+ T cell frequencies in the three sampling sites. Student’s t-test and ANOVA.

### S. aureus PGN reduced gut CD4+ T cells through apoptosis of IgG-bound CD4+ T cells in EcoHIV mice

Furthermore, S. aureus PGN decreased CD3+CD4+ T cells (Figure 3D) and increased surface-bound autoIgG on CD4+ T cells in the gut, but not in blood or spleen (Figure 4A–4D). IgG+CD4+ T cells were highly apoptotic at all sites, in contrast to IgG–CD4+ T cells (Figure 4E-4G). CD4−CD3+ T cell frequencies were unaffected, with minimal IgG binding (Figure S3). In vitro, S. aureus PGN did not directly induce gut CD4+ T cell apoptosis (Figure S4). These results indicate that S. aureus PGN indirectly drives gut CD4+ T cell apoptosis through autoIgG binding.

**Figure 4.**
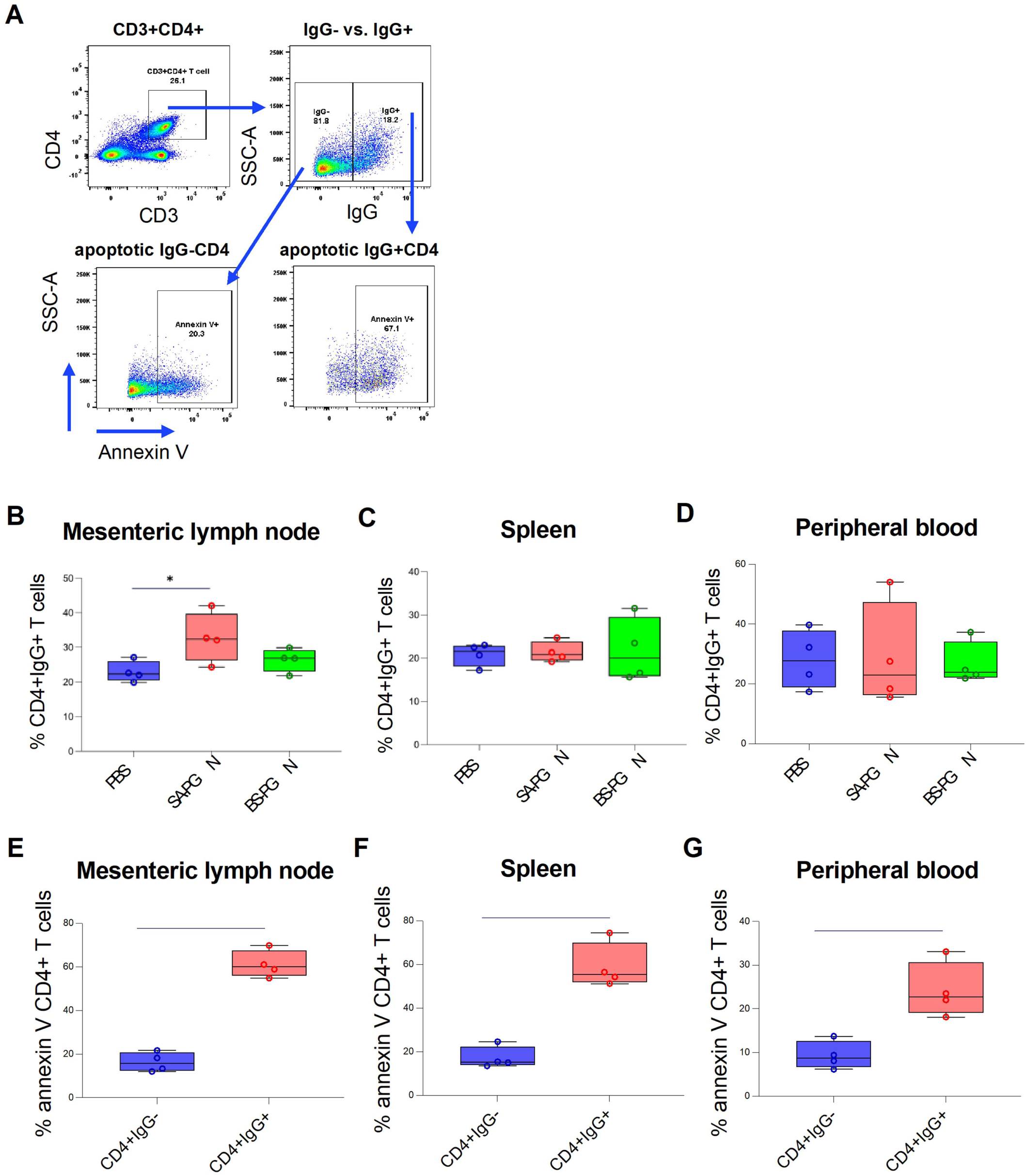
S. aureus PGN increases apoptotic IgG+CD4+ T cells in gut-associated tissues. (A) Annexin V binding on IgG+ vs. IgG– CD4+ T cells. (B–D) Frequencies of IgG+CD4+ T cells in mesenteric lymph nodes, spleen, and blood. (E–G) Annexin V binding on IgG+ vs. IgG– CD4+ T cells. Student’s t-test and ANOVA.

### S. aureus PGN drives class switch DNA recombination (CSR) via TLR2 in vitro

Both native and heat-inactivated S. aureus PGN promoted CSR through TLR2, unlike B. subtilis PGN (Figures 5A–5B, S5A–S5C, S6). Notably, B. subtilis PGN induced stronger TLR2 signaling than S. aureus PGN (Figure S5D), suggesting that S. aureus PGN–mediated CSR involves additional mechanisms beyond TLR2 activation.

**Figure 5.**
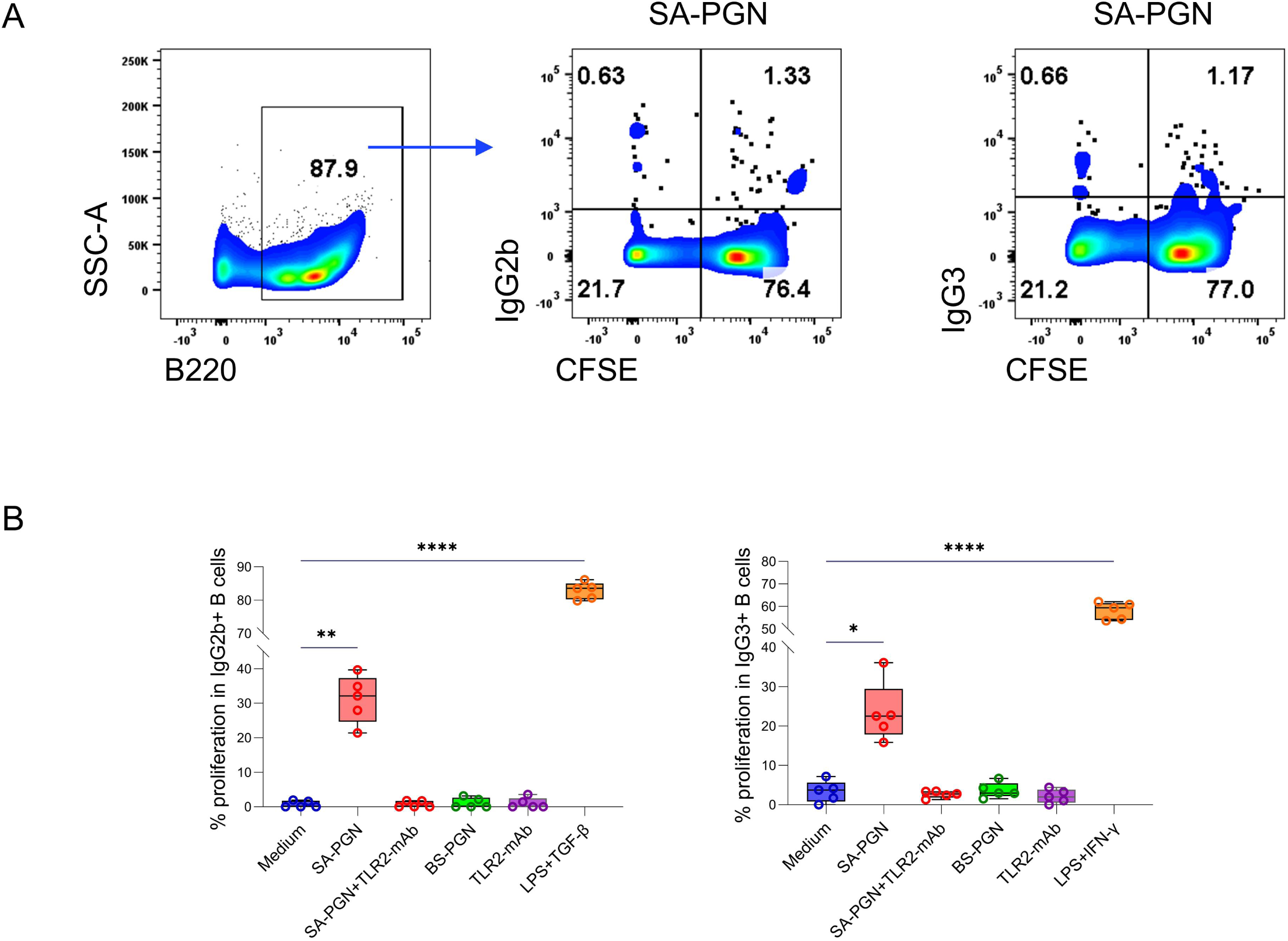
S. aureus PGN induces CSR to IgG2b and IgG3 in vitro. Splenic B cells from naive C57BL/6J mice were cultured with medium, and PGNs (10 μg/mL) ± TLR2-mAb or isotype Ab (1 μg/mL) for 96 h. Percentages of proliferating IgG2b+ and IgG3+ B cells (%CFSE^low^IgG2b/IgG3+ in B220+ cells) are shown (A–B). ANOVA.

### PGN structural analysis

S. aureus PGN (∼18 disaccharide units) contains muramic acid (Mur) and the amino acids Lys, Ala, Gly, and Glu, with a molar ratio of 1.4 Ala: 2.9 Gly: 1.0 Glu: 1.1 Lys (GC/MS). In contrast, B. subtilis PGN (80–250 disaccharide units) contains Mur, Dpm, Ala, and Glu, with a molar ratio of 1.0 Ala: 1.0 Glu: 0.8 Dpm, and includes meso-Dpm and D-Glu enantiomers. Structurally, S. aureus PGN is classified as A3α (L-Lys–Gly₅₋₆, L-Lys–Gly₃), while B. subtilis PGN is A1γ (meso-Dpm–direct), accounting for their distinct immunomodulatory properties.

## Discussion

Autoimmunity contributes to infectious disease pathogenesis, even in the absence of clinical autoimmune disease (1). We first reported anti-CD4 IgG autoimmunity in HIV in 2017, linking it to CD4+ T cell depletion and poor immune recovery post-ART (3), a finding later supported by others (15, 16). Here, we show that S. aureus translocation drives anti-CD4 IgG production, potentially explaining higher CD4+ T cell recovery failure in African PWH with prevalent S. aureus colonization (17).

Anti-CD4 IgG was first described in the pre-ART era (18), but its pathogenicity remained unclear until our 2017 study demonstrated its role in poor CD4+ T cell recovery post-ART (3). Recent studies show that anti-CD4 IgG levels rise during untreated HIV infection, decline with ART, yet persist in a subset of PWH on ART; an inverse correlation with CD4+ T cell counts was observed at 48 and 96 weeks post-ART, but not before ART or at 24 weeks (10), while other reports found no association after prolonged ART (5). Low-affinity, non-cytotoxic natural autoantibodies may emerge from chronic inflammation (19), whereas pathogenic autoantibodies require disordered somatic hypermutation and CSR over time. Unlike pre-ART anti-CD4 IgGs, which are non-cytotoxic natural autoantibodies likely induced by viremia and inflammation, pathogenic anti-CD4 IgGs in PWH on ART mediate ADCC and develop post-ART (15, 20–22). Anti-prothrombin IgG (aPT IgG) is an antiphospholipid antibody directed against prothrombin; our study reveals that plasma levels of anti-prothrombin IgG were directly correlated with anti-CD4 IgG levels, but not with CD4+ T cell counts, suggesting that these two autoantibodies may be generated through a similar mechanism and anti-prothrombin IgG does not contribute to CD4+ T cell counts. Previous studies (23, 24) have shown increased levels of anti-prothrombin IgG in HIV-infected, predominantly ART-untreated individuals; however, these antibodies were typically transient and not strongly associated with thrombotic events or overt antiphospholipid syndrome.

S. aureus colonization and antigen translocation alone may not trigger pathogenic autoantibody production without a compromised barrier, immune perturbations, or genetic predisposition to autoimmunity. Unlike autoantibodies in autoimmune diseases, pathogenic anti-CD4 IgG in PWH on suppressive ART may arise from: 1) elevated apoptotic CD4+ T cells increasing CD4 self-antigen in lymph nodes (25); 2) persistent viral protein release, including HIV gp120, enhancing CD4 binding to anti-CD4 IgG and promoting its production (26); 3) the absence of autoimmune-prone genetic backgrounds in most PWH differentiates these responses from classical autoimmune diseases.

CD4+ T cell apoptosis was detected only in mesenteric lymph nodes, not in the spleen or blood, after S. aureus PGN administration. Although IgG+CD4+ T cells exhibited higher levels of apoptosis than IgG−CD4+ T cells across all sites (Figure 4E-4G), increased IgG surface binding on CD4+ T cells following *S. aureus* PGN intraperitoneal injection was observed only in mesenteric lymph nodes, and not in the spleen or blood (Figure 4B-4D). The site-specific effects likely reflect differences in local exposure to SA-PGN following intraperitoneal administration. Mesenteric lymph nodes are expected to encounter higher concentrations of SA-PGN than the spleen or peripheral blood. Since SIV/HIV infection initiates CD4+ T cell loss in the gut during acute infection and progresses to peripheral depletion in chronic stages (27), we hypothesize that prolonged exposure to S. aureus PGN (e.g., 1 year) may also deplete peripheral CD4+ T cells. Importantly, mesenteric CD4+ T cell death was not induced by PGN directly in vitro, suggesting an indirect mechanism via autoIgG binding and cell apoptosis. The mechanisms driving S. aureus PGN-mediated gut autoIgG production and CD4+ T cell decline require further studies.

Previous studies reveal that bacterial lipoteichoic acid (LTA), a TLR2 ligand, primarily induces TLR2-dependent pro-inflammatory cytokines rather than autoantibodies (6, 28). Generally, TLR2 signaling in B cells more often supports cytokine secretion, activation marker upregulation, or differentiation than direct mitogenesis, comparable to strong mitogens such as CpG (TLR9) or anti-IgM (29). In addition, the downstream pathways and outcomes of TLR2 activation can differ in cytokine production and proliferation. S. aureus PGN exhibited structural differences compared with B. subtilis PGN. An additional mechanism may account for S. aureus PGN-mediated B cell CSR, which warrants further investigation.

The limitations of this study are: 1) there is no figure reported by the PGN structure analysis from the company (Creative Proteomics, Shirley, NY, USA); 2) the potential contamination of protein A or LTA in S. aureus PGN. Nonetheless, several lines of evidence support PGN as a driver of autoantibody production: (1) a TLR2 inhibitor blocked PGN effects; (2) PGN, but not the other Gram+ bacterial products, is detectable in human circulation without a clinical bacterial infection (8); (3) S. aureus protein A and enterotoxin B reduce lupus autoimmunity in mice (30, 31); and (5) LTA, a TLR2 ligand, primarily induces pro-inflammatory cytokines rather than autoantibodies (6, 28).

Currently, no treatment is available for improving CD4+ T cell recovery in PWH on suppressive ART. Up to 20% of PWH on ART fail to restore peripheral CD4+ T cell counts to the levels of healthy controls; increased morbidity and mortality have been demonstrated in these patients, which is the main challenge in the HIV clinic. This study reveals that *S. aureus* and its PGN translocation may drive anti-CD4 autoimmunity and hinder immune recovery in PWH on suppressive ART, highlighting *S. aureus* colonization as a therapeutic target and supporting the development of competitive probiotic interventions such as *Bacillus subtilis* (32).

## Materials and methods

### Human subjects

This study enrolled 32 controls (without HIV), 32 HIV+/ART-naïve, and 53 HIV+/ART+ individuals (>2 years undetectable plasma HIV-1 RNA) from Sahlgrenska University Hospital (Gothenburg, Sweden), San Francisco General Hospital (San Francisco, CA, USA), Ralph H. Johnson Veteran Health Care System (RHJ VAHCS, SC, USA), and the Medical University of South Carolina (MUSC, SC, USA). The local institutional review board approved the study. Demographics and clinical details are provided in Table 1. PWH were divided into low and high anti-CD4 IgG subgroups using a 35 ng/ml cutoff, based on the 95^th^ percentile in controls.

**Table 1.**
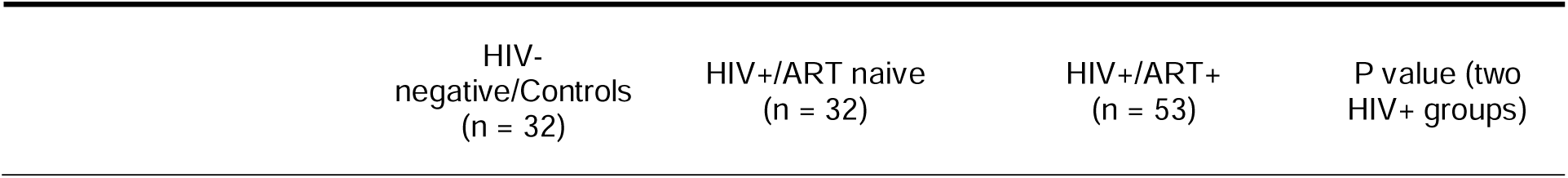
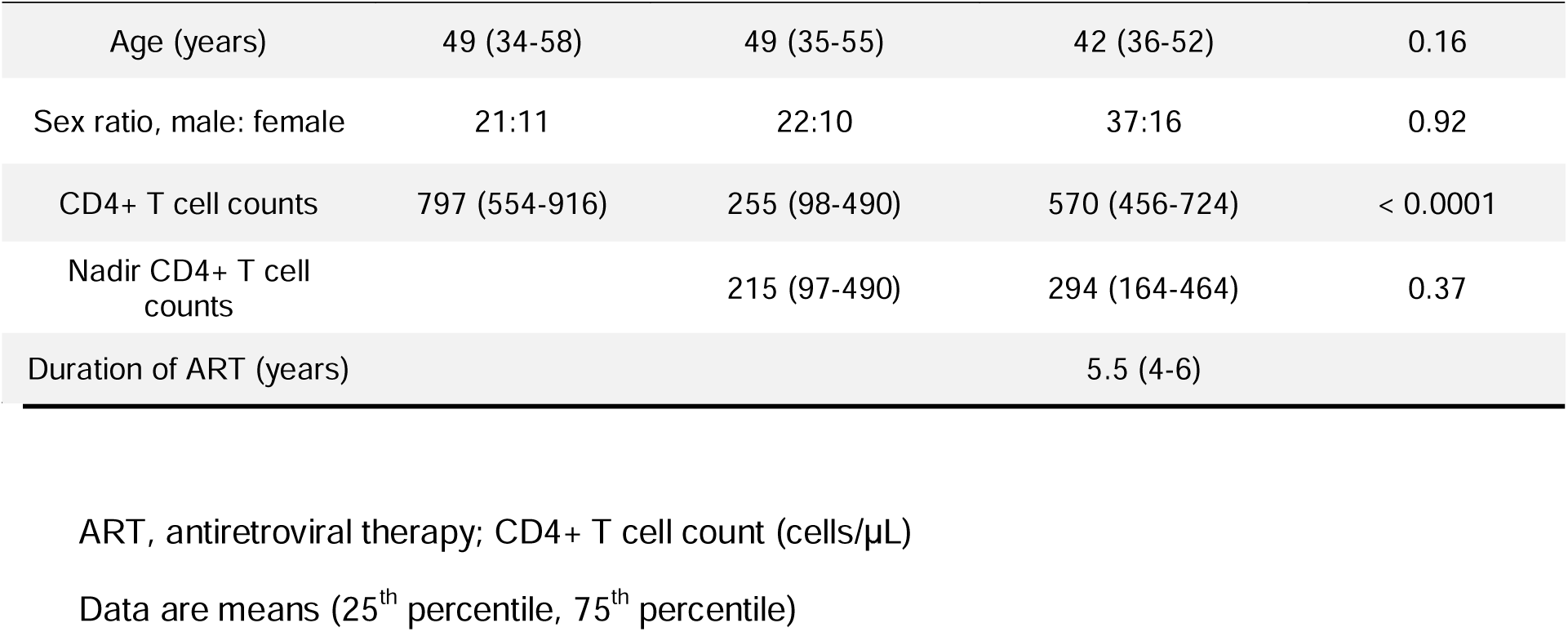
Clinical characteristics of study participants.

| HIV-<br>negative/Controls<br>(n = 32) | HIV+/ART naive<br>(n = 32) | HIV+/ART+<br>(n = 53) | P value (two<br>HIV+ groups) |
| --- | --- | --- | --- |

|  |  |  |  |  |
| --- | --- | --- | --- | --- |
| Age (years) | 49 (34-58) | 49 (35-55) | 42 (36-52) | 0.16 |
| Sex ratio, male: female | 21:11 | 22:10 | 37:16 | 0.92 |
| CD4+ T cell counts | 797 (554-916) | 255 (98-490) | 570 (456-724) | < 0.0001 |
| Nadir CD4+ T cell counts |  | 215 (97-490) | 294 (164-464) | 0.37 |
| Duration of ART (years) |  |  | 5.5 (4-6) |  |

ART, antiretroviral therapy; CD4+ T cell count (cells/ μL)
Data are means (25<sup>th</sup> percentile, 75<sup>th</sup> percentile)

### Autoantigen protein array, B-cell gene profiles, and plasma levels of anti-CD4 autoantibodies

We reported the methods of protein array containing 87 self-antigens, B cell gene microarray, and plasma anti-CD4 autoantibody levels (3, 7).

### Plasma *S. aureus* antigen translocation

*S. aureus* lysate was prepared using the BugBuster Master Mix and coated onto high-binding plates (Sino Biological, Wayne, PA) at 2 μg/100 μL/well. The plate was blocked with 3% BSA and added with diluted plasma samples (1:4000 for IgA, 1:400000 for IgG). Detection was performed using horseradish peroxidase-labeled anti-human IgG and IgA.

### Bacterial PGN structural analysis

PGNs from *S. aureus* (ATCC, 6538P) and *B. subtilis* (NR-607, BEI, Manassas, VA) were separated with an Agilent HPLC system and analyzed using mass spectrometry (Creative Proteomics, Shirley, NY, USA).

### Mice Experiment

Mouse study design was shown in Figure 3A. EcoHIV plasmid was obtained from Dr. Hu (Virginia Commonwealth University) (33). Six-week-old female C57BL/6J mice (Jackson Laboratory, Bar Harbor, ME) received mixed antibiotics (Gibco™, 1 mg/mL) in drinking water for 3 days and a 2-day break. Mice were injected i.p. with EcoHIV (50 μL, 3.45 × 10^8^ IU/mL; n = 6/group), followed by i.p. of PBS, S. aureus PGN, or B. subtilis PGN (InvivoGen, San Diego, CA) twice weekly for 12 weeks. Whole blood was drawn via the tail vein method from EcoHIV-injected mice under aseptic conditions. To verify ecoHIV infection, the blood was used to isolate DNA via the Quick-DNA/RNA Miniprep Plus Kit (Zymo; Irvine, CA, Cat # D7003). The standard curve was generated from viral titer and threshold cycles using the ABM qPCR lentivirus titer kit (LV900). qPCR for HIV-LTR was performed for the unknown and standard samples via QuantaBio PerfecTa Toughmix (QuantaBio, Beverly, MA), supplemented with the following primers and a probe: LTRF, 5’-CTGGCTAATTAGGGAACCCACTG-3’; LTRR, 5’-GGACTAAACGGATCTGAGGGATCTC-3’; and the LTRP, 5’-(FAM)-TTACCAGAGTCACACAACAGACGGGCA-(MGBNFQ)-3’. The thermal program used was 3 min at 55°C, 10 min at 95°C, and 40 cycles of 15 sec at 95°C and 1 min at 60°C. qPCR was performed at the Bio-Rad CFX96 qPCR instrument (Bio-Rad Laboratories, Hercules, CA), and data analysis was done with the built-in Bio-Rad software. The number of proviral copies in the unknown sample is the number of the total number of copies minus the number of copies in the control reaction. All samples were run in triplicate.

### Mouse serum anti-CD4 IgG levels

Serum anti-CD4 IgG was detected using recombinant mouse CD4 protein (1 μg/100 μL/well) and horseradish peroxidase-labeled anti-mouse IgG or its subclasses.

### B cell class switch DNA recombination (CSR)

B cells, isolated from spleens of 8-week-old mice (STEMCELL B Cell Negative Isolation Kit), were cultured with medium, S. aureus PGN, or B. subtilis PGN (10 μg/mL), ± anti-TLR2 or isotype IgG2a (100 ng/mL, InvivoGen), or positive controls (LPS from E. coli 055:B5 [3 μg/mL, InvivoGen] plus IFN-γ [50 ng/mL, BioLegend] or TGF-β1 [2 ng/mL, R&D Systems]). CSR was measured by proliferating IgG3 and IgG2b B cell percentages using a BD FACSVerse™ Cell Analyzer after staining with anti-B220, anti-IgG2b, anti-IgG3, anti-CD138, and anti-IgM antibodies (BioLegend).

### IgG surface binding on CD4+ T cells

Cells from blood, spleen, and mesenteric lymph nodes were stained with anti-CD3, anti-CD4, anti-IgG, Annexin V, and streptavidin, then analyzed on a BD FACSVerse™ flow cytometer using FlowJo software.

### PGN-mediated TLR2 activity

TLR2 activity was detected using the HEK-Blue™ mTLR2 cells (InvivoGen) following the manufacturer’s instruction.

### Statistics

Data were analyzed using Student’s t-test or Mann–Whitney test for two-group comparisons and one-way ANOVA for multiple groups. Associations were assessed by Spearman correlation. A P value < 0.05 was considered significant.

## Supporting information

all figures

## Authors’ contributions

D.C. wrote the first version of the manuscript. D.C., Z.L., T.S., D.J., and W.N. performed experiments and analyzed data. M.G., R.W., S.H., and Z.L. recruited the study participants. D.C., Z.L., W.J., L.C.N., R.P., R.H., W. H., M.G., JEM, and R.A. were involved in critically revising the manuscript.

## Acknowledgments

We thank John D. Dinolfo, Ph.D. (MUSC), for English language editing. This work was supported by the Ralph H. Johnson VA Health Care System I01CX002422 (Jiang) from the U.S. Department of Veterans Affairs; the VA Southeast Network 7 Research Development Award (Adekunle); NIH grants R01DA059854 (Jiang), R01DA059538 (Jiang), R01DA056876 (Hu), P30AI027767 (Saag/Health), R01NS094067 (Price), UL1TR001450, TL1TR002382 (Adekunle), and UL1TR002378 (Adekunle, TL1). The content is solely the responsibility of the authors and does not necessarily represent the official views of the NIH.

## Availability of data and materials

Data are available.

## Figure legends

**Figure S1.**
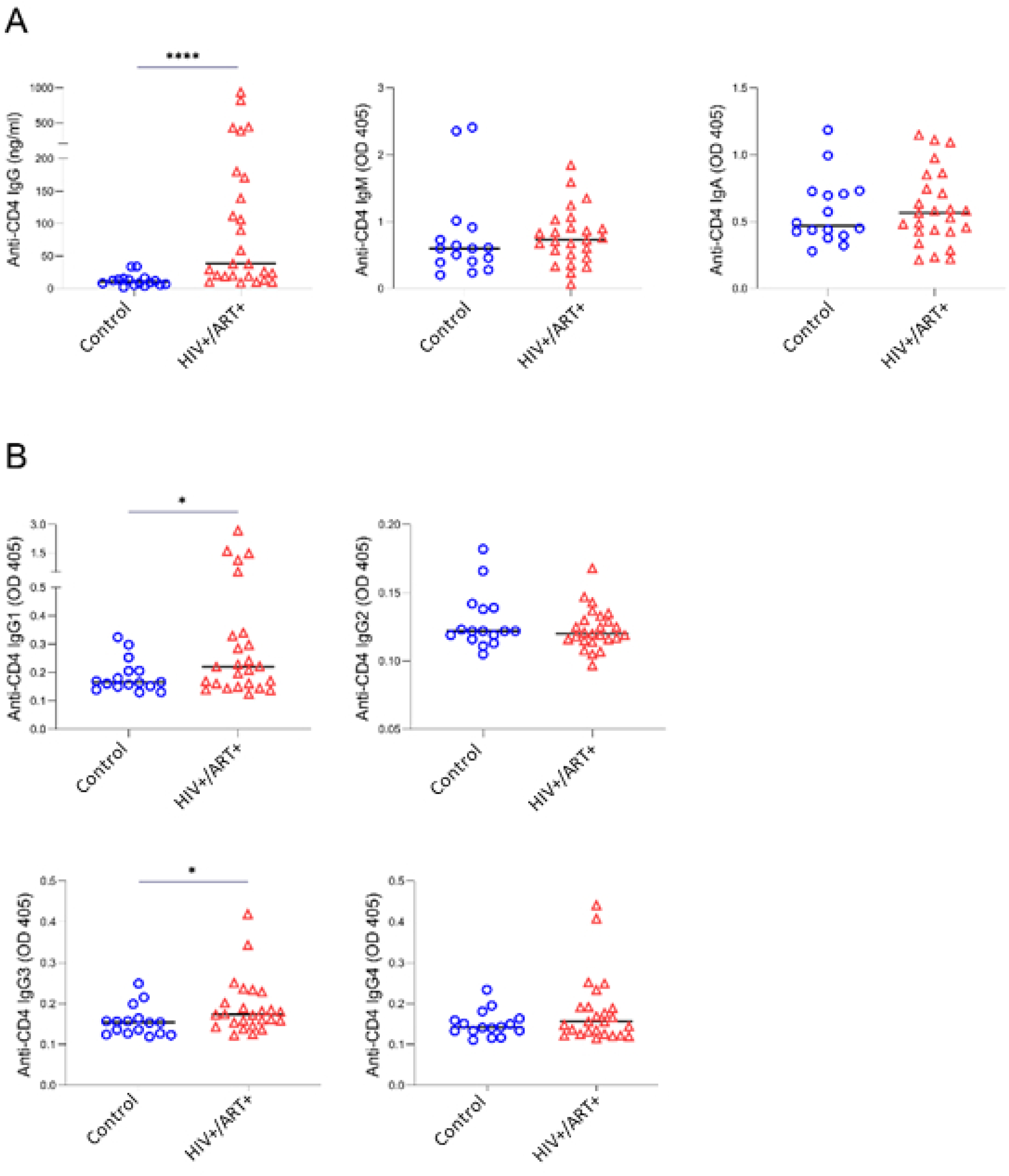
Elevated plasma anti-CD4 IgG1 and IgG3 subclasses in PWH on ART. (A) The median plasma levels of anti-CD4 IgG, IgM, and IgA in controls (n = 15) and HIV+/ART+ subjects (n = 23). (B) The median plasma levels of anti-CD4 IgG subclasses (IgG1-IgG4) between the two groups. Mann-Whitney tests.

**Figure S2.**
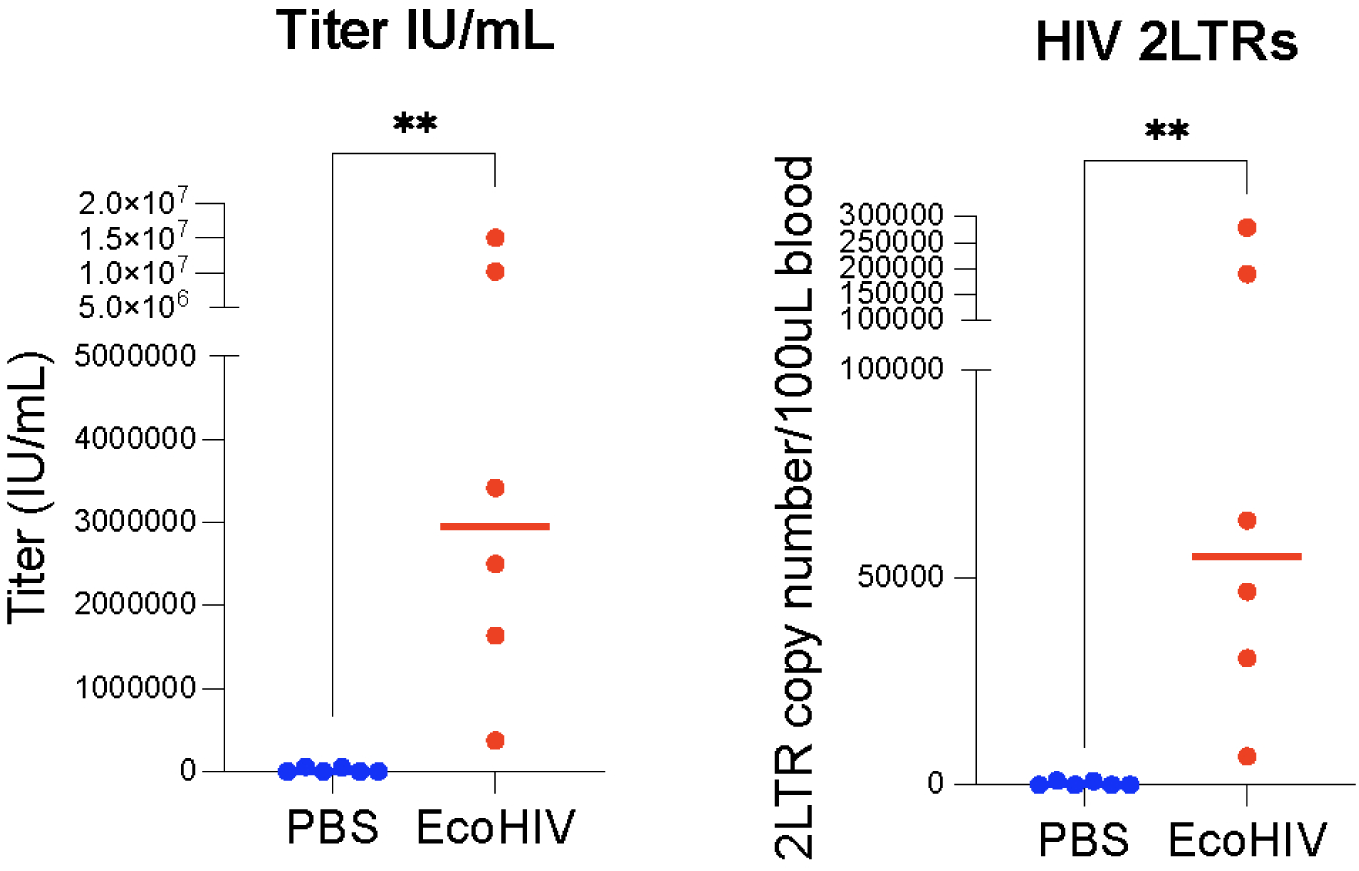
Confirmation of established EcoHIV infection in mice. The EcoHIV virus was verified by qPCR in blood samples collected one month after infection.

**Figure S3.**
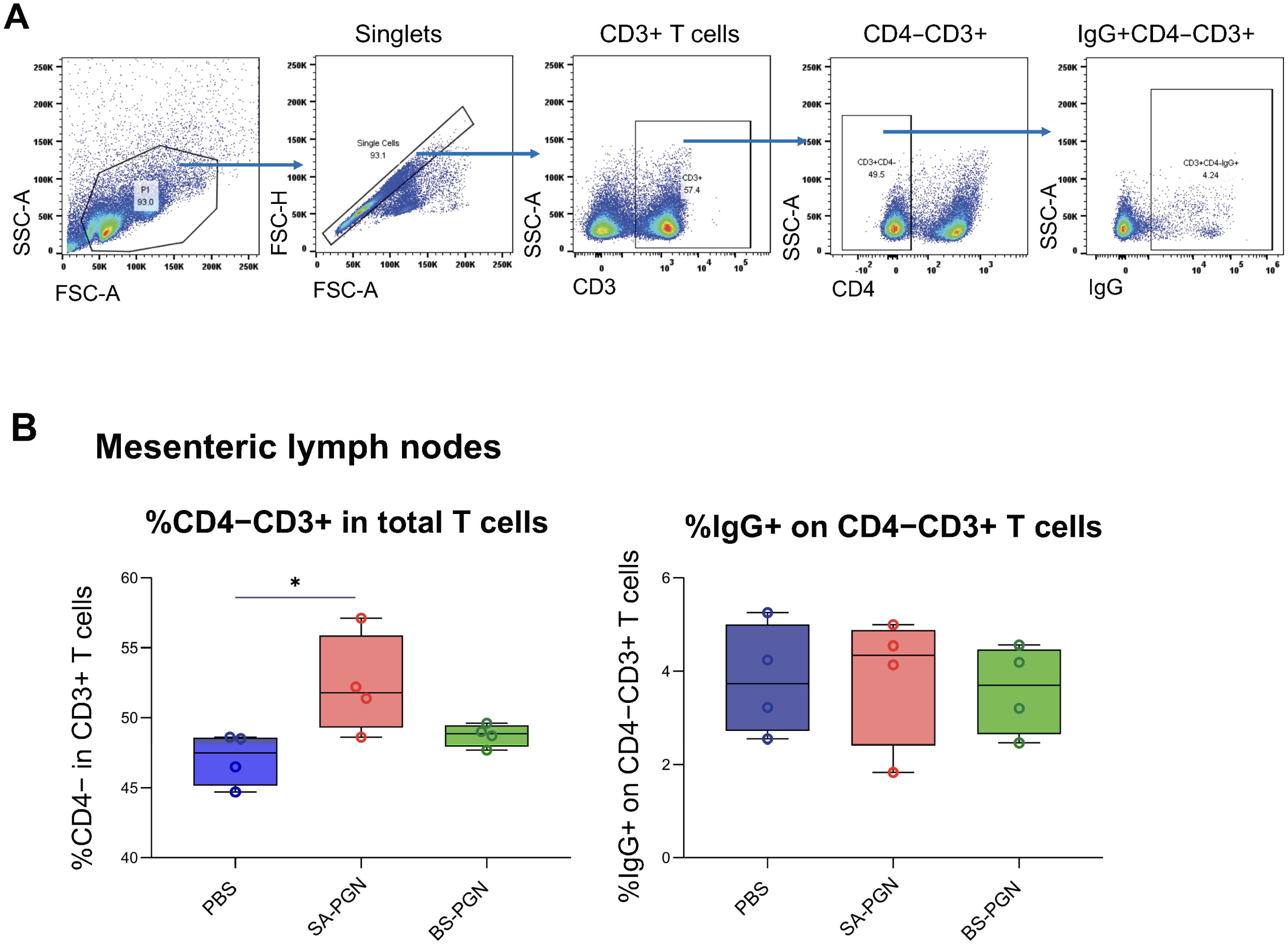
S. aureus PGN does not affect gut CD4−CD3+ T cells in vivo. (A) Gating strategy. (B) Percentages of CD4−CD3+ T cells among total CD3+ T cells and IgG+CD4−CD3+ T cells in mesenteric lymph nodes. ANOVA.

**Figure S4.**
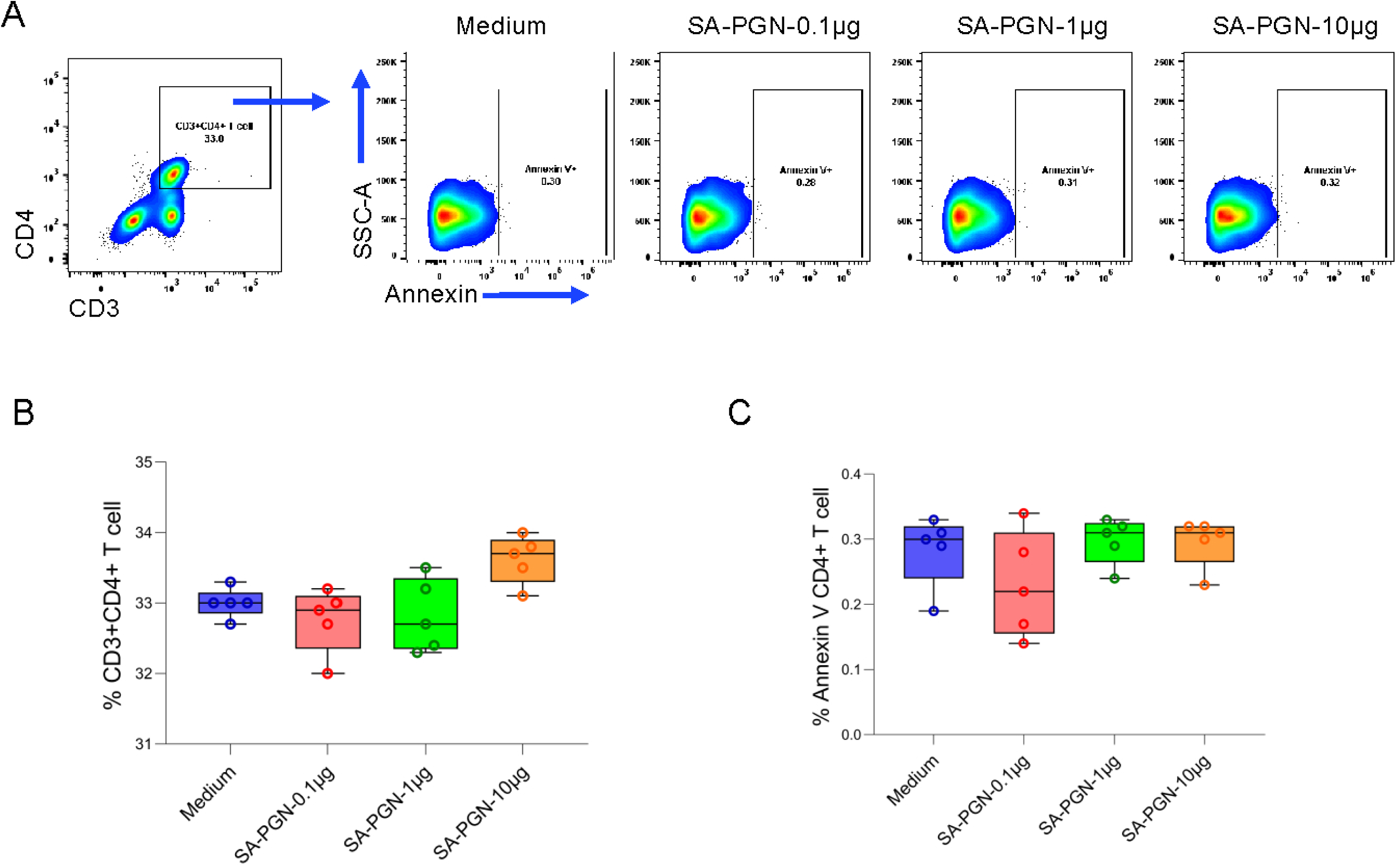
S. aureus PGN does not directly induce gut CD4+ T cell apoptosis in vitro. Lymphocytes from mesenteric lymph nodes were treated with S. aureus PGN (0.1–10 μg/mL) for 4 days. (A-B) Percentages of CD4+ T cells and Annexin V+CD4+ T cells. ANOVA.

**Figure S5.**
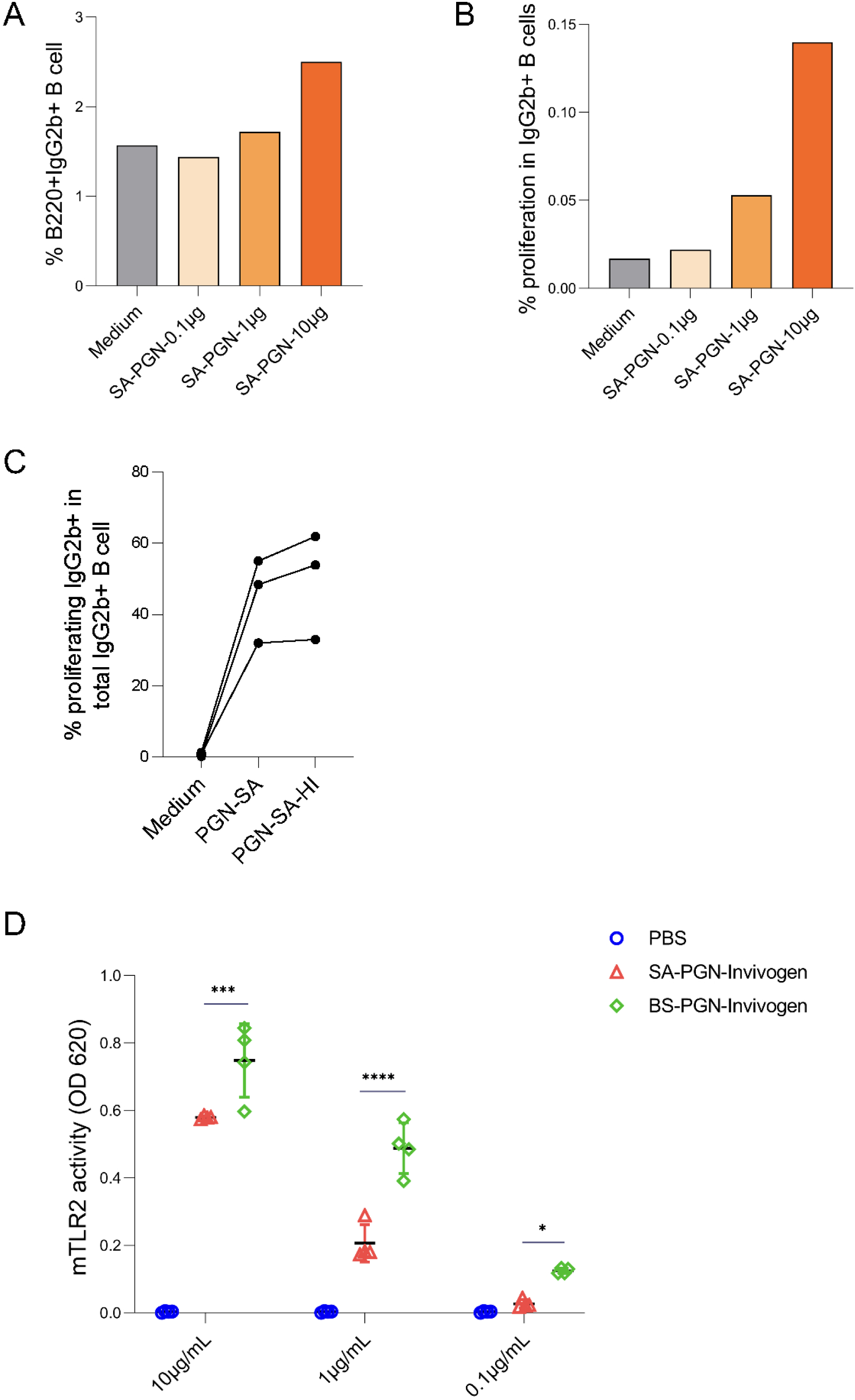
PGN dose titration for CSR and TLR2 activity. (A–B) S. aureus PGN induced total (A) and proliferating (B) IgG2b+ B cells. (C) CSR induction by heat-inactivated vs. native S. aureus PGN. (D) mTLR2 reporter cell responses after 16 h treatment. ANOVA.

**Figure S6.**
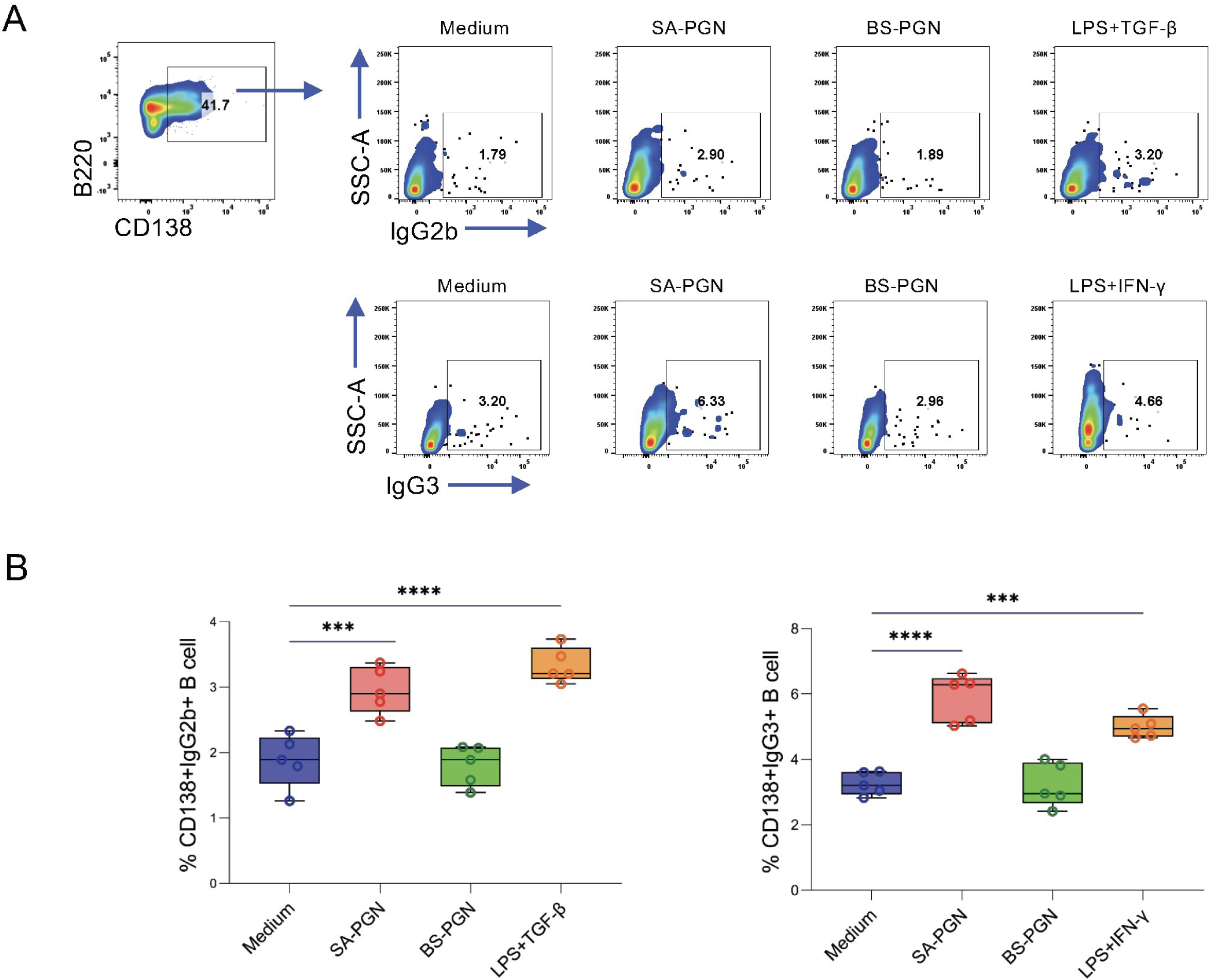
S. aureus PGN induces CSR in vitro. Splenic B cells from naive mice were cultured with PGN (10 μg/mL) for 96 h. Percentages of CD138+ IgG2b+ and IgG3+ plasmablasts (%CD138+CFSE^lowIgG2b/IgG3+ in B220+ cells) are shown (A–B). ANOVA.

